# Spatiotemporal multiplexing on subretinal prosthesis increases selectivity of the network-mediated stimulation

**DOI:** 10.64898/2026.11.11.766021

**Authors:** Nathan Jensen, Keith Ly, Davis Pham-Howard, Mohajeet Bhuckory, Daniel Palanker

## Abstract

Subretinal photovoltaic implants (PRIMA) provide form vision in patients blinded by age-related macular degeneration. The success of this approach is attributed in part to the network-mediated stimulation, which preserves many features of the natural retinal signal encoding. For efficient stimulation of bipolar cells (BC) with smaller pixels, electric field confinement should be relaxed, yet direct activation of the retinal ganglion cells (RGC) should be avoided. Here we present a new method for selective stimulation of bipolar cells using image multiplexing – sequential presentation of the image’s components in several subframes. With sizes of the subframe islands optimized by computational modelling, the selectivity of network (BC) mediated retinal stimulation increased over 5 times in rats. Using this approach, electric field confinement can be relaxed, allowing pixels as small as 20 µm to efficiently stimulate bipolar cells, while avoiding direct stimulation of RGCs and preserving high contrast of the pattern. This software-based improvement enables higher visual acuity of prosthetic vision without a change in implanted hardware.

## 1 Introduction

Microelectrode arrays (MEA) have emerged as reliable bi-directional interfaces with neurons. In read-out applications, they have been utilized for providing artificial limbs and voice using brain signals-driven actuators and speakers. MEA-mediated stimulation has further been used in treating Parkinson’s tremor, paralysis, in auditory and visual prostheses, and in other applications [1–8]. Many of the first neural interfaces targeted large groups of neurons, but advances in decoding the brain’s activity [9] and in fabricating high-density microelectrode arrays [10] have allowed the recording and activation of neurons approaching single-cell resolution. Accordingly, neural selectivity – the ability to selectively stimulate different types of neurons – has grown in importance.

The fidelity of brain-machine interfaces is often limited by their ability to target smaller groups of specific neurons. For example, pitch of the electrode array in the subretinal prosthesis PRIMA limits the resolution of prosthetic vision: clinical trials demonstrated acuity closely matching its 100 µm pixel pitch [11], while in rodents, this trend continues with smaller pixels down to the species’ natural limit of 28 µm resolution [12]. The ability of the PRIMA implant to provide high-fidelity form vision is attributed in part to its targeting of retinal bipolar cells (BC) thereby preserving much of the natural downstream signal processing in the retinal neural network [13–16]. In fact, stimulating other cells in the retina, such as retinal ganglion cells (RGC) and their axons, caused distorted percepts [17, 18] and dramatically reduced resolution [19]. In the retina, where different neurons are uniquely stratified into layers, neural selectivity can be achieved by proper shaping of electric field.

Tight confinement of electric field from local return electrodes increases the stimulation threshold, especially with smaller pixels. Relaxing the field confinement – such as by using monopolar electrodes [20] – reduces the thresholds but results in the excessive stimulation of RGCs. Balancing these two considerations limits the pixel pitch to no smaller than 40 µm (corresponding to a visual acuity of 20/160), even when employing 3-D electrodes [21].

Here we present a method to overcome such limitations by using spatiotemporal multiplexing of the images projected onto the stimulating array. Similar approaches have been employed to reduce MEA crosstalk [22, 23], and here we demonstrate its additional capability to shape the vertical penetration of electric field. With a monopolar MEA, the penetration depth of an electric field in tissue is related to how many neighboring electrodes are simultaneously activated. Therefore, by splitting a single video frame into several sequentially activated sub-images, one can dramatically increase the neural selectivity of subretinal prostheses. Critically, such image multiplexing does not increase the stimulation thresholds of the nearby bipolar cells, but dramatically decreases the activation of more remote neurons – the RGCs.

Using computational modeling and ex-vivo recordings with a custom fabricated 22 µm pitch monopolar MEA, we demonstrate very significant changes in the vertical extent of electric field with image multiplexing. We also confirm this effect by electrophysiological recordings in Royal College of Surgeon (RCS) rats using synaptic blockers to isolate the direct stimulation of RGCs. Multiplexed images increased the RGC threshold by 5.4 times on average, compared to simultaneous activation of the full field, without affecting the BC thresholds. This software-based approach provides a path to high-acuity prosthetic vision in human patients without a need to change the implants’ hardware.

## 2 Results

For a monopolar high-density MEA, the penetration depth of the electric field into tissue is defined by the collective size of the current-injecting area. Approximating the illuminated zone as a disk electrode of radius *a* emitting current density *j*, its electric potential in the medium with resistivity *ρ* scales with distance *z* approximately as 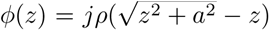 [24]. Near the electrode, at *z ≪ a*, electric field is constant (*E*(*z*) = *jρ*), and electric potential decreases linearly with distance. At distance *z* = *a*, it drops by about half, and at *z* ≫ *a*, it decreases as *ϕ*(*z*) ≈ 1*/z*, and *E*(*z*) ≈ 1*/z*^2^. Since the field penetration depth is similar to the size of the activated zone on the array *a*, limiting the illuminated area attenuates the electric field *E* in deeper layers of the retina, while remaining constant near the electrode. In RCS rats, the targeted bipolar cells are roughly 50 µm in height, with dendritic terminals residing directly above the subretinal implant. The off-target neurons (RGCs) reside in the ganglion cell layer, and their axons in the nerve fiber layer (NFL), which is roughly 80 µm above from the implant. Therefore, with an activated area of about 100 µm in size, bipolar cells should experience nearly the same field as with a larger illuminated zone at constant current density *j*, while RGCs should experience a weaker field than with wide-field stimulation.

Therefore, to control the penetration depth of electric field into tissue, the full image can be broken into islands illuminated sequentially within the time of one frame of the video stream, as illustrated in Fig. 1. A multiplexed image is presented as a sequence of its intersections with a few (four in the illustrated example) subsets of such islands, which we call subframes. For a steady perception of a pulsed video stream, the frame rate *F* is typically at or above 30 Hz, which allows for flicker fusion to occur [25]. The minimum frame duration is the inverse of the frame rate, *T* ≤ 1*/F*, and for *F* = 30 Hz, *T* ≈ 33 ms. With 4 subframes, illustrated in Fig. 1, the image of each frame is broken into complimentary patterns exposed during a subframe time *t* = 8 ms. With 3 subframes, *t* could be extended to 33*/*3 = 11 ms.

**Fig. 1.**
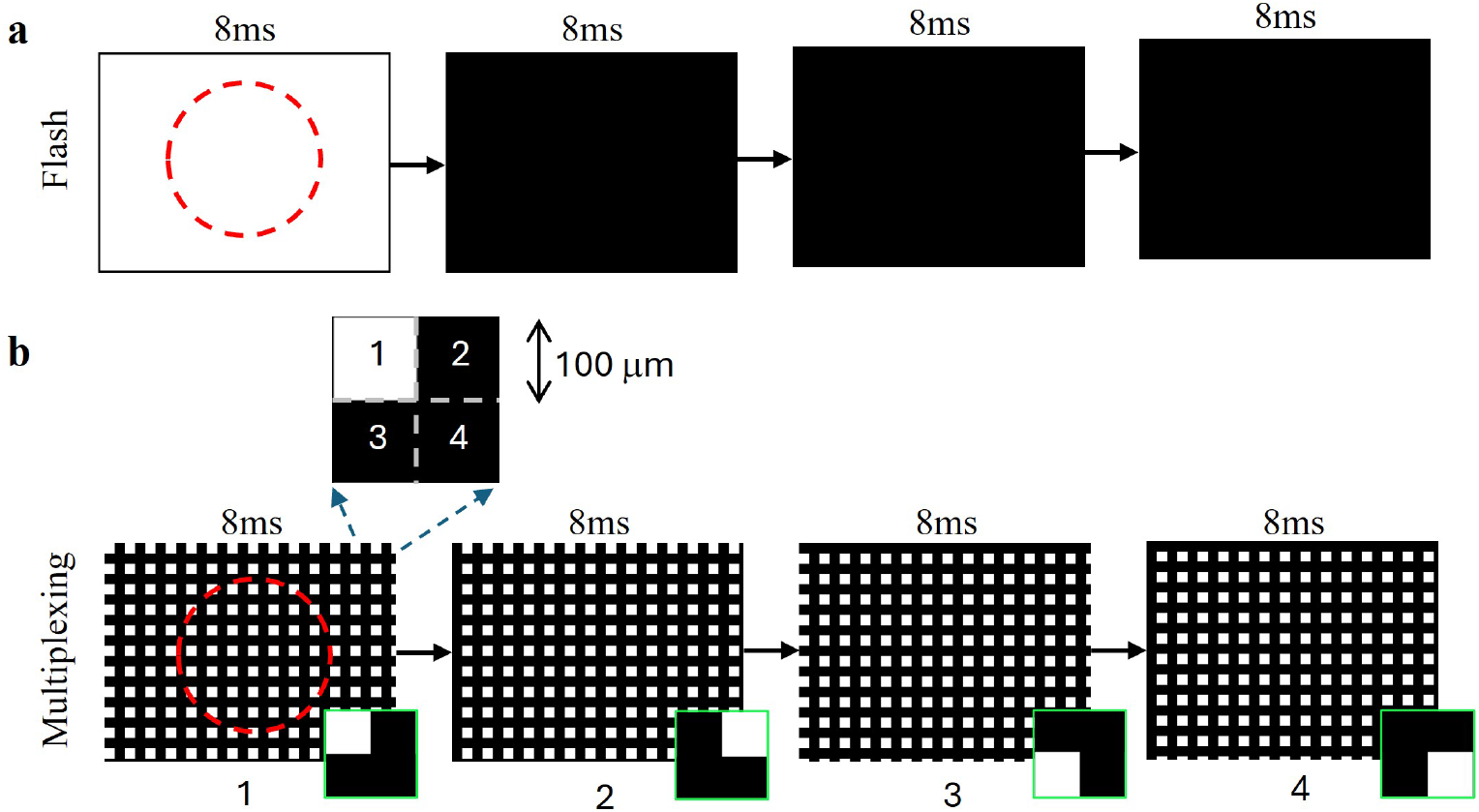
Stimulation protocols for a full-field flash and multiplexing. **a**) A single frame of full-field illumination for 8 ms followed by 3 frames of no stimulation of 8 ms each. **b**) In multiplexing protocol, stimulation consisted of 4 frames at 8 ms each, composed of 100 µm squares in different quadrants (see inset). Red dashed circle indicates outline of the implant.

### 2.1 Modeling the electric field in the retina

To assess the effects of image multiplexing, we simulated the electric field generated by the MEA in rat retina. The MEA used in this study is a photovoltaic monopolar array with a global return electrode located on the backside of the device [26]. Each pixel in the array consists of a photodiode, a shunt resistor, and an active electrode. The electrode-electrolyte interface for 8 ms pulses at a 2 Hz repetition rate is represented by a simplified Randles Circuit composed of the double layer capacitance and access resistance of the electrode. All electrodes are coupled via common electrolyte, which is modeled as an electrical conductance matrix. Electric field in the tissue was calculated by combining the described circuit with elementary electric fields calculated using finite element methods in COMSOL Multiphysics [20, 27, 28]. We employed Neumann boundary condition (uniform current density) on the active electrodes since their RC time constant of 2.7 ms is below the stimulus duration of 8 ms. On the global return, we used Dirichlet (equipotential) boundary condition since its RC time constant of 420 ms is much larger than the stimulus duration. The RC time constant serves as an estimate of the transition time between the initial equipotential state and the ultimate steady state of constant current density [29].

The photocurrent generated by each pixel is defined by light intensity of the projected image. We compared a full-field image (the entire implant is illuminated), and the 4× multiplexed image, where each subframe is composed of square tiles (e.g. 100x100 µm shown in Fig. 1). The neural response of a bipolar cell can be closely approximated by the drop in electric potential between its axonal and dendritic terminals [30]. In RCS rats, this corresponds to 0um (dendritic end) and 52um (axonal end) above the implant. In human retina (PRIMA patients), BCs are the same length but start roughly 35 µm above the implant [25, 31]. Similarly, the axonal activating function of RGCs can be approximated as the drop in electric potential along the axons [21], which reside in the NFL (82 µm above the implant in RCS rats and 120 µm in patients).

To find the optimum island size, we sampled them from 25 to 1200 µm, with the largest size being equivalent to a full-field illumination. The electric field across the BCs and at the NFL plane from a subset of these simulations is shown in Fig. 2a. The electric potential for on-target (BCs) and off-target (RGCs) neural stimulation is plotted vs. the size of the islands in Fig. 2b. With multiplexing, the neural selectivity ratio (BC/RGC) is maximized by using islands of 100 µm in width. Compared to the full-field image, this slightly increases BC stimulation and significantly decreases the RGC stimulation. As a result, the ratio of BC stimulation strength to that of RGCs increased 5.7-fold, as shown in Fig. 2c.

**Fig. 2.**
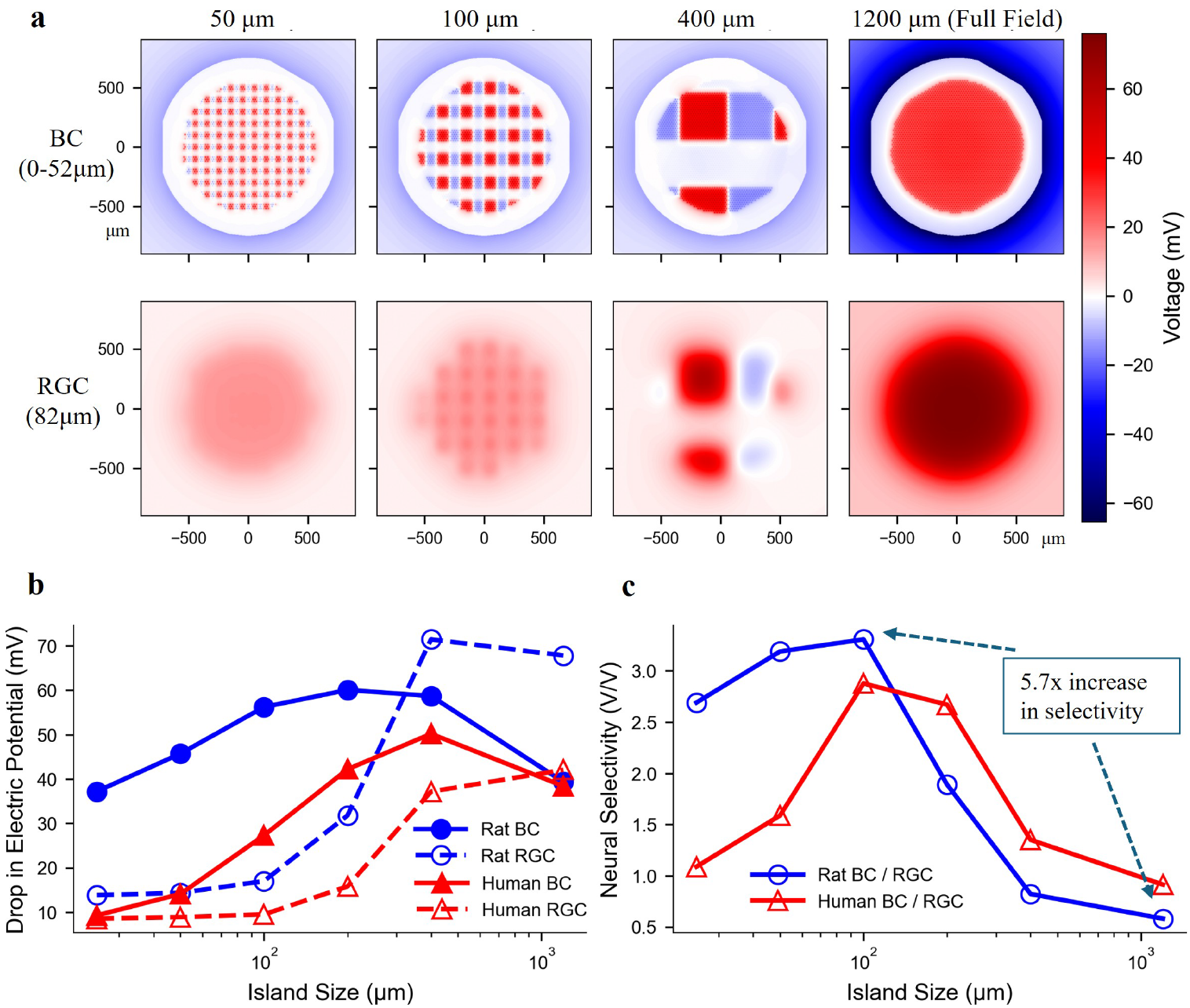
Simulations of electric field in the retina. **a**) The average electric field across bipolar cells (BC, top) and retinal ganglion cells (RGC, bottom) in RCS rats, during 1 out of 4 subframes with multiplexing islands of various sizes. The largest size (1200 µm) corresponds to the entire implant being illuminated during one subframe, equivalent to the full-field single flash stimulus. **b**) The maximum drop in electric potential across the BC’s and in RGC plane plotted vs. island size, for rat and human retinas. **c**) The BC/RGC neural selectivity ratio for rats and humans as a function of island size. The ratio peaks around 100 µm for both species. The model predicts the neural selectivity to increase 5.7 times by using multiplexing, compared to the full-field flash.

### 2.2 Measurement of electric field in electrolyte

To validate computational modeling, we recorded the electric potential generated by the photovoltaic array in Hank’s balanced salt solution (HBSS) diluted to mimic retinal resistivity (1000 Ω·cm). Illumination was provided using a microscope-mounted digital micromirror device (DMD). The recording pipette electrode was positioned 5 µm, 57 µm (for BCs), and 82 µm (for RGCs) above the implant (Fig. 3a). We could not record below 5 µm due to occasional crashing of the pipette walls. Uniform electrolyte in a Petri dish is a rough approximation of the retina in the eye, with expected deviations in the electric potential compared to physiologically accurate simulations of a thin retina under highly conductive vitreous.

**Fig. 3.**
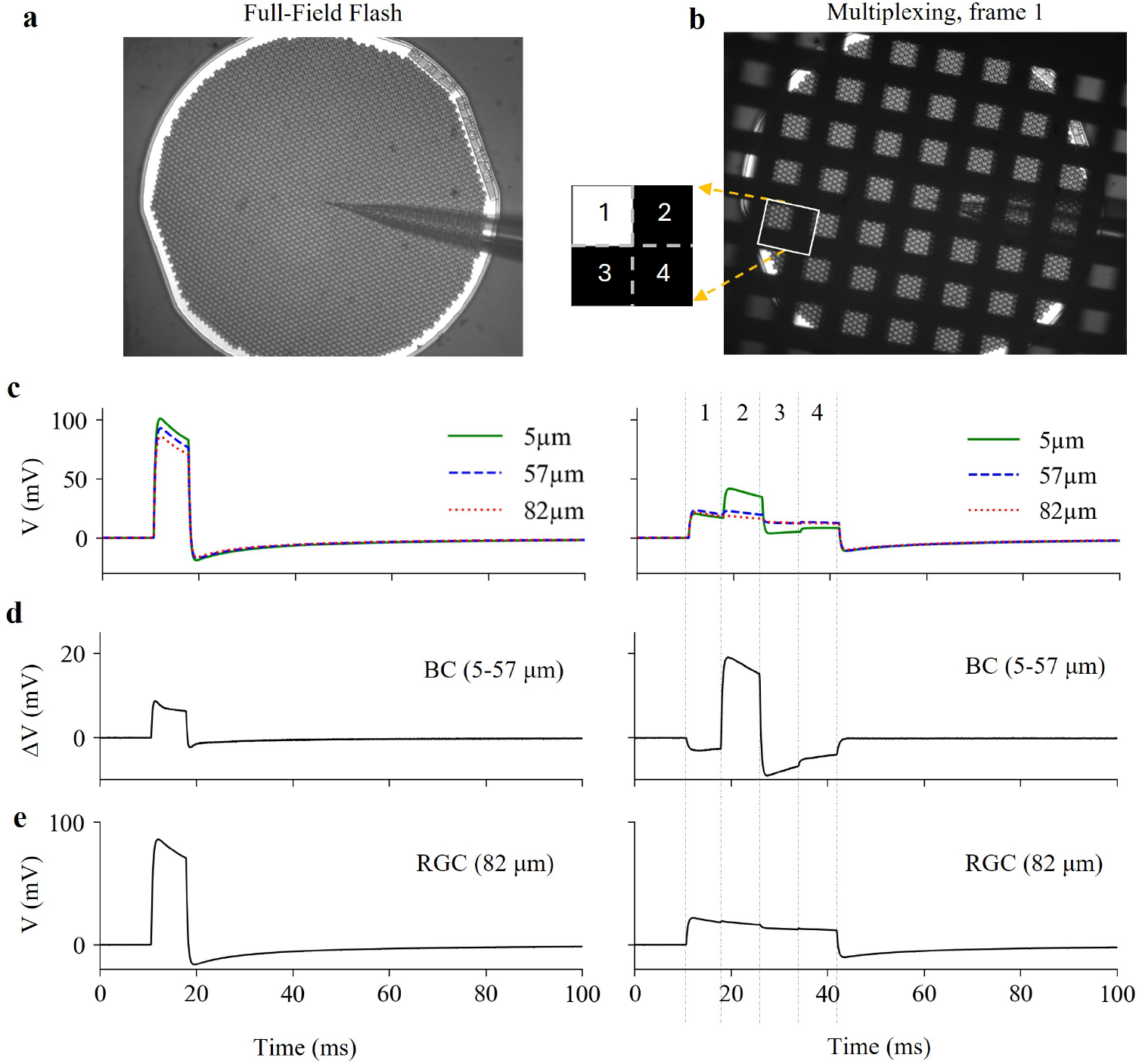
Recordings of electric potential in saline. **a**) Recording pipette is positioned above the array, illuminated by a full-field flash for 8 ms **b**) One of 4 subframes composed of 100 µm square islands. **c**) Electric potential in saline 5, 57, and 82 µm above the implant with a full-filed (left) and multiplexed (right) illumination. **d**) Stimulation strength of BCs, estimated by subtracting the electric potential at 57 µm above the implant from that at 5 µm. **e**) Stimulation strength of RGCs, estimated as the electric potential in the NFL - at 82 µm above the implant.

Pulsed near-infrared (NIR) illumination was projected at a 2 Hz frame rate, using either a single image during the entire frame of 8 ms, or 4 subframes consisting of tiled square patterns, each presented for 8 ms in sequence (Fig. 3a,b). The recorded signals are shown in Fig. 3c. The potential difference between the top and bottom of BCs is shown in Fig. 3d, and for RGCs – in Fig. 3e. As a result of multiplexing (right column in Fig. 3), the voltage drop across the BC planes increased by 2.2*×*, while the maximum electric potential in the NFL plane decreased by a factor of 3.9. Since stimulation thresholds in terms of irradiance change reciprocal to the measured electric potential, the spatiotemporal multiplexing is expected to significantly increase the selectivity of network-mediated retinal stimulation (RGC/BC thresholds ratio) compared to a single-frame full-field illumination.

### 2.3 VEP thresholds in rodents

RCS rats were implanted with subretinal photovoltaic MEAs of 22 µm electrode pitch. A single 8 ms flash and a multiplexed version of a full-field illumination were projected onto the retina at 2 Hz repetition rate (Fig. 4b). VEP was recorded via transcranial electrodes placed above the visual cortex, and thresholds were found by extrapolation to the noise level. The corneal signal was also recorded to subtract the stimulus artifact from the VEP waveforms (example waveforms shown in Fig. 4a). Intravitreal injection of the neurotransmitter blockers was applied to isolate the direct stimulation of RGCs.

**Fig. 4.**
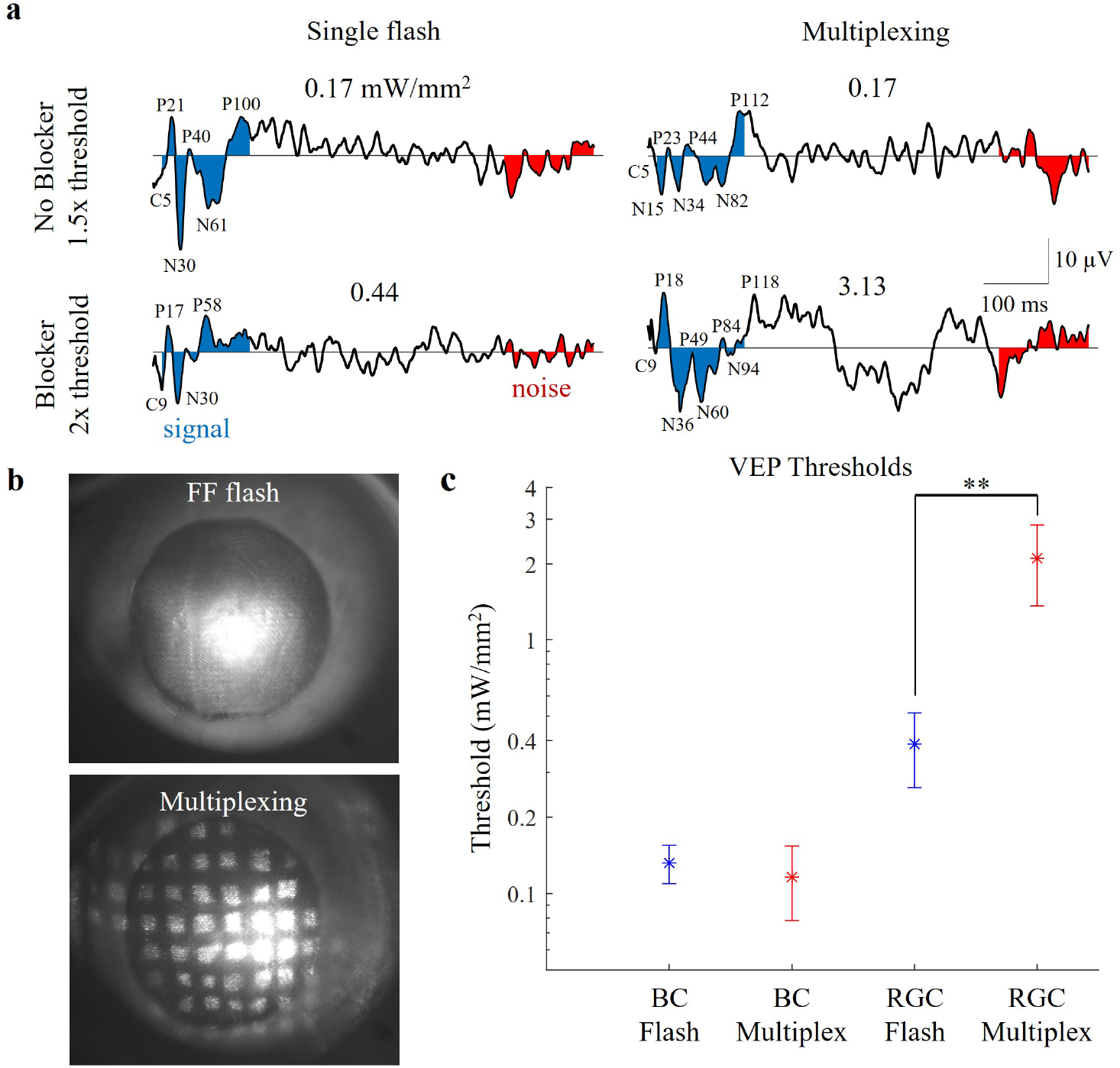
Cortical responses and thresholds with full-field flashes and multiplexing. **a**) VEP waveforms at the same irradiance ratio above threshold (1.5*×* for no blocker, 2*×* for blocker), for a single full-field flash and a multiplexed image. C labels the first trough, P for peaks, and N for all following troughs, with the number following each letter as the time in milliseconds for each feature. Typical responses to single flashes are faster and have fewer features, whilst multiplexing responses are longer and have significantly more features due to longer stimulus. **b**) Fundus images with a full-field illumination above and multiplexed (one subframe shown) illumination below. **c**) VEP thresholds with a single flash (n = 5) and multiplexed image (n = 5) for both BCs (no blockers) and RGCs (with blockers). For a flash, the BC threshold was 0.13*±*0.02 mW/mm^2^ and RGC threshold - 0.39*±*0.12 mW/mm^2^, resulting in 3*×* selectivity. For the multiplexed image, the BC threshold was 0.12*±*0.04 mW/mm^2^, and the RGC threshold - 2.11*±*0.74 mW/mm^2^, resulting in 17*×* selectivity, demonstrating 5.9*×* improvement with multiplexing.

The stimulation threshold of bipolar cells was 0.13±0.02 and 0.12±0.04 mW/mm^2^ for FF and its multiplexed version, respectively. The slight decrease in BC threshold with multiplexing, expected from the model, was not statistically significant in these measurements. Following the intravitreal injection of synaptic blockers, which prevent synaptic transmission from all cells to the RGCs, the thresholds increased to 0.39±0.12 mW/mm^2^ for a single flash and to 2.11±0.74 mW/mm^2^ for multiplexing. When using blockers, the threshold with multiplexing increased by 5.9× more than with the FF illumination, which is statistically significant (*p* = 0.0037).

### 2.4 Predicted selectivity in human retina with bipolar pillar arrays

Subretinal implants will perform better in human patients with carefully designed geometry, such as bipolar arrays with pillar electrodes which were optimized by analyzing the electrical stimulus strength and contrast of Landolt C patterns [21]. In such devices, the laser irradiance had to be reduced to prevent activation of RGC axons, which limited their stimulation strength.

To predict the performance of such arrays with multiplexing in human patients, we tested it on the same tiled Landolt C pattern used in [21] (Fig. 5a), with gap sizes ranging from 24 to 120 µm. Bipolar cells were placed 37–57 µm above the implant and RGC axons at 120–140 µm, as seen in human anatomy [31]. For the multiplexed version of the pattern, 150 µm wide islands were used to segment the image into 4 subframes. These islands are larger than the 100 µm optimum found for rats because the BCs and the NFL reside farther from the implant in human patients.

**Fig. 5.**
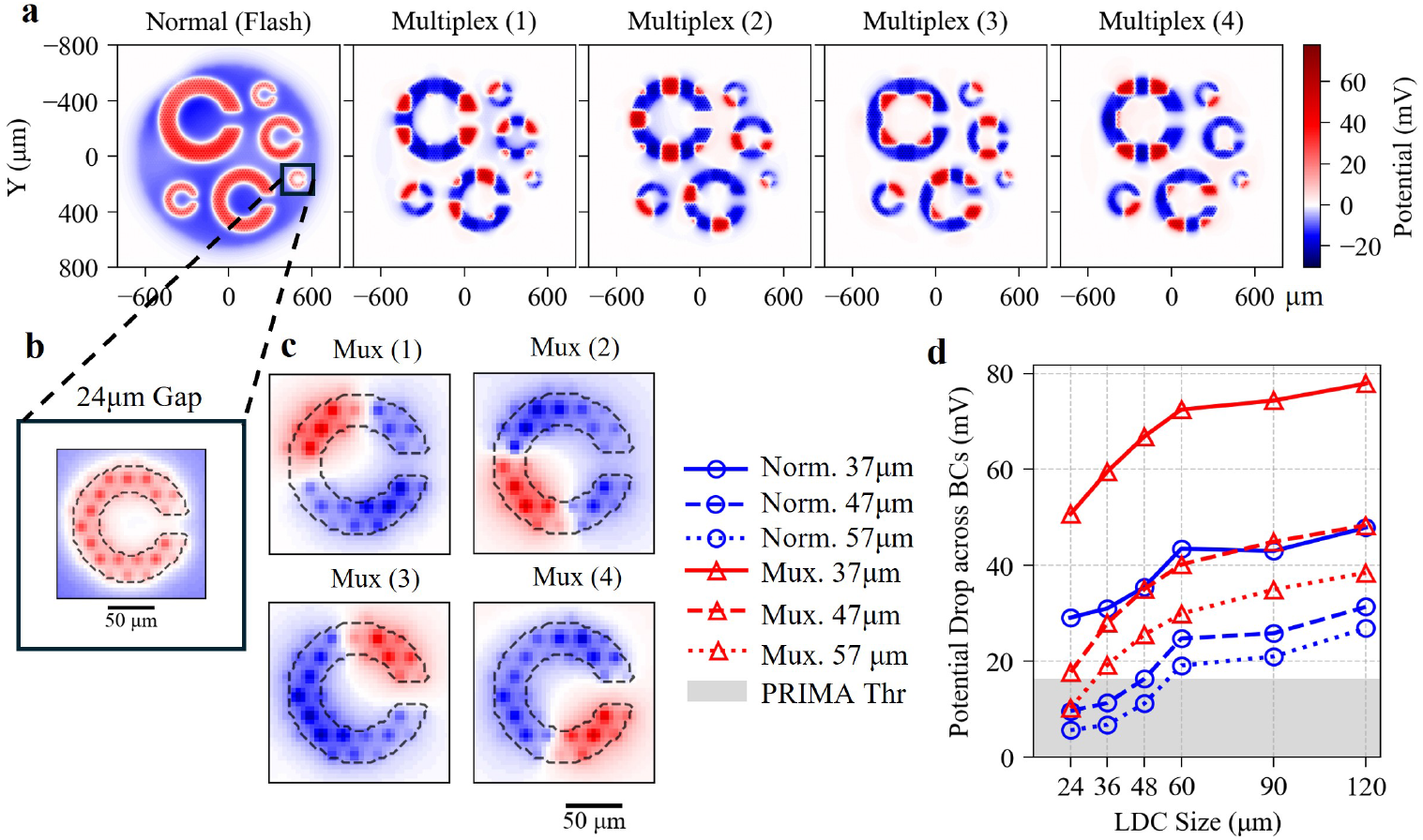
Simulation of electric field with bipolar pixels and pillar electrodes in human retina. **a**) The time-averaged electric potential across BCs with dendritic ends 37 µm above the implant, for the normal flash where the entire image is projected at once for 8 ms, and for the 4 subframes of the multiplexed image. **b**) Zoom of the smallest Landolt C with a 24 µm gap for the normal projection, where the entire letter is illuminated simultaneously. **c**) The smallest Landolt C for the multiplexing (Mux) case, illuminated in a sequence of 4 subframes. **d**) Maximum electric potential for each size of Landolt C gap, for the normal flash stimulus (blue, circles), and the Multiplexed (Mux) stimulus (red, triangles). Three different heights of a debris layer above the implant were simulated to predict performance for a variety of patients, 37 µm (solid), 47 µm (dash), and 57 µm (dotted). The irradiances were 2.2 mW/mm^2^ without multiplexing and 3.0 mW/mm^2^ with, to keep RGC stimulation strength below 20 mV. Shaded region indicates the worst case clinical threshold recorded in PRIMA patients of 16.4 mV.

The attenuated field near RGCs with multiplexed illumination allows increasing the laser irradiance up to the 3.0 mW/mm^2^ used in clinical trials [11], while keeping the drop of electric potential in the NFL plane below 20 mV (as with PRIMA implants). Additionally, multiplexing lowers the fill factor of the implant (the fraction of pixels illuminated at any given time), thus weakening the negative electric potential over non-illuminated areas. Both factors increase the stimulation strength, which was the limiting factor of optimized devices under normal illumination [21]. For example, with 37 µm debris, the BC stimulation strength of the smallest Landolt C (24 µm) increased from 29 to 51 mV. In fact, multiplexing provides stimulation strength above the human stimulation threshold for the smallest Landolt C in all but the thickest debris (Fig. 5d).

Contrast of the letter, however, depends on the islands’ placement: for the smallest Landolt C, contrast was only 78% when the entire letter was contained in one subframe (e.g. Fig. 5b), but 92% when the subframes split it equally into 4 sections, as shown in Fig. 5c.

## 3 Discussion

We demonstrated that spatiotemporal multiplexing of images projected onto photovoltaic subretinal prostheses increases the neural selectivity of retinal stimulation. Computational modeling predicted a 5.7-fold increase in the ratio of RGC to BC thresholds, and benchtop measurements confirmed an expected change in neural selectivity. In RCS rats, multiplexing increased the RGC threshold 5.4 times, and the selectivity ratio by 5.9 times – in close agreement with the model. This occurs because the penetration depth of the electric field scales with the size of the illuminated area, and therefore subframes of smaller islands reduce the electric field at the NFL, without affecting the near field stimulating the BCs.

Balancing neural selectivity against stimulation threshold previously limited the pixel pitch of subretinal implants to no smaller than 40 µm [21]. Since multiplexing decreases the RGC stimulation without increasing the BC thresholds, it relaxes this trade-off and may enable pixels as small as 20 µm in patients, corresponding to a visual acuity of about 20/80. Multiplexing also comes with the benefit of reducing the fill factor of the implant, preventing large images from having a significantly increased stimulation threshold with bipolar arrays as shown in [21]. Our simulations predict that these benefits carry over to bipolar pillar arrays in human retina, although these predictions remain to be confirmed clinically.

Furthermore, the image processing methods used here allow for tuning the electric field to individual patients by software control. The height of the inner nuclear layer (INL) and NFL above the implant can be found using optical coherence tomography (OCT) imaging, and an optimal island size can then be selected based on these measurements (e.g. Fig. 2). This is important because the thickness of a subretinal debris layer varies significantly among patients [31], which is likely to affect the efficacy of stimulation [21]. The island sizes can even be adjusted at different regions of the implant, to account for variations in retinal thickness within a single patient, although this would require eye tracking.

It was previously demonstrated that crosstalk between electrodes in an MEA can be reduced using transient return electrodes, including in subretinal prostheses [12, 22, 32]. Multiplexing inherently provides the benefits of transient returns since electrodes of the illuminated pixels charge-up in each subframe, and better conduct the return current during the subsequent subframes. Therefore, this scheme can improve contrast in addition to the neural selectivity, but contrast optimization for features smaller than the islands will require image-dependent subframe partitioning and assignment. In this study, VEP thresholds were recorded at 2 Hz frame rate. We assume that with multiplexing at 30 Hz frame rate, the flicker fusion will be as complete as with single frames at the same repetition rate. Techniques similar to multiplexing have been successfully used in other neural interfaces, such as continuous interleaved sampling in cochlear implants [33], nevertheless the perceptual effects of multiplexing should be validated in clinical tests.

Another limitation of this study is that ex-vivo measurements were performed in a uniform medium, and the model approximates the retinal resistivity as uniform across its thickness, whereas the actual tissue is layered and heterogeneous. In vivo, only the full-field illumination was compared to multiplexing, and its effect on sparser patterns remains to be assessed, especially with human patients.

By increasing the selectivity of network-mediated stimulation without raising the bipolar cell thresholds, spatiotemporal multiplexing offers a path toward high-acuity prosthetic vision. It can be implemented using software control, without any changes to hardware of subretinal implants. Image multiplexing could also be used for optimizing the electric field confinement with dense pixel arrays in other neural interfaces, such as cortical implants, though additional studies would be required on a case-by-case basis.

## 4 Methods

### 4.1 Implants

All animals were implanted with monopolar photovoltaic MEAs of 30 µm in thickness, 1.5 mm in diameter, composed of 22 µm wide hexagonal pixels with 12 µm in diameter active electrodes in their centers. Each pixel has a single photodiode and a shunt resistor connected between the active and return electrodes, which allows the electrodes to discharge between pulses. All return electrodes in the pixels are connected to one global return electrode, located on the back side of the device. Active and return electrodes were both coated with sputtered iridium oxide film. The fabrication process of these implants and their characterization was described in detail previously [26].

### 4.2 Modelling of electric fields in rodents

We modeled the electric field generated in the retina by such subretinal arrays. The electrode-electrolyte interface for 8 ms pulses at 2 Hz repetition rate is well approximated by a simplified Randles circuit consisting only of the double layer capacitance and access resistance of the electrode [34, 35]. Electric coupling between electrodes in a shared electrolyte was represented by an electrical conductance matrix, as described in [20]. The photocurrent generated by each pixel was determined based on the projected image, either a full-field illumination or a 4 spatiotemporally multiplexed image, where each subframe was composed of square tiles ranging from 25 µm to 400 µm. We utilized Xyce-based RPSim to solve this circuit model, obtaining individual currents at each electrode.

To compute the electric field in tissue, the electrode currents were averaged during each pulse (normal) or subframe (multiplexing) and used as coefficients to linearly combine a set of elementary electric fields, which were generated by the finite element method (FEM) in COMSOL Multiphysics to solve the Poisson equation of volume conduction. We employed Neumann boundary condition (uniform current density) on the active electrodes since their RC time constant of 2.7 ms is below the stimulus duration of 8 ms. On the global return, we used Dirichlet (equipotential) boundary condition since its RC time constant of 420 ms is much larger than the stimulus duration [29].

The retina was modelled as a 160 µm thick homogeneous tissue with resistivity 1000 Ω·cm. To approximate the neural response in Royal College of Surgeons (RCS) rats, the resulting electric potentials were evaluated at heights of 0 µm and 50 µm above the implant, which correspond to the dendritic and axonal terminals of bipolar cells, respectively. Activating function of the retinal ganglion cells (RGCs) was evaluated as the drop in electric potential 82 µm above the implant, where the nerve fiber layer (NFL) resides.

### 4.3 Modelling of neural stimulation in human retina

We modeled the electric field in the retina generated by a subretinal bipolar photovoltaic array with 20 µm pixel pitch, having pillar electrodes described in [21]. In this array, each pixel consists of a photodiode, a shunt resistor, an active electrode on top of a 35 µm tall pillar, and a mesh of local return electrodes around each pixel. The electrode-electrolyte interface was modelled similar to as was described in section 4.2, but with Neumann boundary conditions used on both the active and return electrodes. Additionally the conductance matrix was simplified following the methods described in [28] with a thresholding ratio of 0.025 and 950 compensation modes. The illumination consisted of tiled Landolt C’s with gap sizes spanning from 24 to 120 µm, with laser irradiance of 3 mW/mm^2^, frame rate of 30 Hz, and pulse width of 9.8 ms. For the multiplexed version of the pattern, 150 µm wide squares were used to segment the image into 4 subframes, and each subframe had a pulse width of 8.3 ms. The electric potential was found in the same manner described in section 4.2, and scaled such that the largest drop in electric potential along the axons at their lowest height (120 µm) was below 20 mV. This resulted in equivalent irradiances of 2.2 mW/mm^2^ without multiplexing and 3.0 mW/mm^2^ with. The electric potential was then extracted across bipolar cells of 52 µm in length, with their dendritic end located at heights varying from 37 to 57 µm above the implant, as observed in human patients [31]. The RGC axons were placed from 120 to 140 µm above the implant, and we ensured that the drop of electric potential in that plane remained below 20 mV, following the procedure used in [21].

### 4.4 Ex-vivo device characterization

For ex-vivo validation of the electric field model, we placed the photovoltaic device in a Petri dish filled with diluted Hank’s balanced salt solution (HBSS) to mimic the 1000 Ω·cm resistivity of retinal tissue, validated by a resistivity probe (HM Digital COM-100). Pulsed near-infrared (NIR) illumination (880 nm) was projected onto the device at a 2 Hz frame rate via an upright microscope. Spatiotemporal distribution of the NIR light was controlled by a digital micromirror device (DMD, Texas Instruments DLP650LNIR), projecting either a single full-field image (8 ms) or its 4× multiplexed version with square tiles of 100 µm (8 ms per sub-frame, 32 ms total).

A recording micropipette was positioned using a micromanipulator (Sutter MPC-200) at vertical distances of 5 µm and 57 µm above the implant to approximate the bipolar cell layer in RCS retina, and at 82 µm to measure the maximum electric potential in the plane where the NFL resides. The electric potential was measured against a Ag*|*AgCl reference electrode, and the signal was averaged over 100 recordings.

### 4.5 Surgical Procedures

All experimental and surgical procedures were conducted in accordance with the Statement for the Use of Animals in Ophthalmic and Vision research of the Association for Research in Vision and Ophthalmology (ARVO) and approved by the Stanford Administrative Panel on Laboratory Animal Care (APLAC). Animals were maintained at the Stanford Animal Facility under 12 h light/12 h dark cycles with food and water ad libitum.

RCS rats between the age of P180 and P300, when photoreceptors were completely lost, were anesthetized with a mixture of ketamine (75 mg/kg) and xylazine (5 mg/kg) injected intraperitoneally. Rats were implanted with three transcranial screw electrodes: one over each hemisphere of the visual cortex (4 mm lateral from midline, 6 mm caudal to bregma), and a reference electrode (2 mm left of midline and 2 mm anterior to bregma). Monopolar photovoltaic devices of 1.5 mm in diameter and 30 µm in thickness with 22 µm pixels were implanted subretinally, as in [36]. Retinal implants were placed in the temporal-dorsal region, approximately 1 mm away from the optic nerve. Complete loss of photoreceptors by P180 and implant position were checked with optical coherence tomography (OCT; HRA2-Spectralis; Heidelberg Engineering, Heidelberg, Germany).

Synaptic blockers (0.8 mM L-AP7, 0.8 mM L-AP4, 0.2 mM NBQX, 0.4 mM strychnine, and 0.32 mM picrotoxin) were injected into the intravitreal space to block the synaptic transmission and isolate RGCs and thereby determine the RGC stimulation thresholds.

To determine the height of BCs and RGCs above the implant, OCT images were taken prior to each recording. Retinal thickness was found to be 79.1*±*14.2 µm.

### 4.6 Electrophysiology

Photovoltaic devices were illuminated with 880 nm NIR laser (M1F-880 nm-400 µm, DILAS, Tucson, AZ) at irradiance ranging from 0.06 to 3.8 mW/mm^2^ with 8 ms pulse duration at 2 Hz repetition rate. Digital micromirror display (DMD; DLP Light Commander, LOGIC PD, Carlsbad, CA) was used for pattern formation, and the image projection optical assembly was mounted on a slit lamp for alignment onto the implant in rat eye, as previously described [12]. Patterns consisted of full-field illumination and a multiplexed image broken into 4 sub-frames (Fig. 4b).

VEPs were recorded using the Espion E3 system (Diagnosys LLC, Lowell, MA) at a sampling rate of 4 kHz and averaged over 250 trials. Rats were anesthetized lightly with subcutaneous injections to ensure a response to patterns. The signal to noise ratio (SNR) of the VEP waveform was calculated by taking the integral of the square of the waveform for 120 ms after the stimulus (signal), divided by the same procedure for the last 120 ms of the waveform (noise) (Fig. 4a). 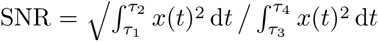 where *x* is the amplitude of the waveform at any time *t*, *τ*_1_ is the start of the signal at 8 ms, *τ*_2_ is the end of the signal at 128 ms, *τ*_3_ is the start of the noise at 380 ms, and *τ*_4_ is the end of the noise at 500 ms.

A paired t-test was used to compare the full-field thresholds to their multiplexed counterparts for each animal (n=5) using MATLAB’s built-in *ttest* function.

## Acknowledgements

Studies funded by National Institutes of Health (R01-EY-035227 and P30-EY-026877); U.S. Department of Defense (W81XWH-22-1-0933); Air Force Office of Scientific Research (AFOSR, FA9550-24-1-0138).

## Author Contributions

N.J. Experiment design, modeling, data analysis, drafting and revising the manuscript. K.L. Experiment design, surgery, electrophysiology, data analysis, drafting and revising the manuscript. D.P-H. Animal care, surgery, electrophysiology. M.B. Supervisory, revising the manuscript. D.P. Supervisory, drafting and revising the manuscript.

## References

[1] Hochberg, L. R. et al. Reach and grasp by people with tetraplegia using a neurally controlled robotic arm. Nature 485, 372–375 (2012). URL 10.1038/nature11076.

[2] Willett, F. R. et al. A high-performance speech neuroprosthesis. Nature 620, 1031–1036 (2023). URL 10.1038/s41586-023-06377-x.

[3] Jia, Q. et al. Wireless closed-loop deep brain stimulation using microelectrode array probes. Journal of Zhejiang University-SCIENCE B 25, 803–823 (2024). URL 10.1631/jzus.B2300400.

[4] Swann, N. C. et al. Adaptive deep brain stimulation for Parkinson’s disease using motor cortex sensing. Journal of Neural Engineering 15, 046006 (2018). URL 10.1088/1741-2552/aabc9b.

[5] Lorach, H. et al. Walking naturally after spinal cord injury using a brain– spine interface. Nature 618, 126–133 (2023). URL 10.1038/s41586-023-06094-5.

[6] Wilson, B. S. & Dorman, M. F. Cochlear implants: a remarkable past and a brilliant future. Hearing Research 242, 3–21 (2008). URL 10.1016/j.heares.2008.06.005.

[7] Holz, F. G. et al. Subretinal photovoltaic implant to restore vision in geographic atrophy due to amd. New England Journal of Medicine (2025). URL https://www.nejm.org/doi/full/10.1056/NEJMoa2501396.

[8] Petrini, F. M. et al. Sensory feedback restoration in leg amputees improves walking speed, metabolic cost and phantom pain. Nature Medicine 25, 1356–1363 (2019). URL 10.1038/s41591-019-0567-3.

[9] Obien, M. E. J., Deligkaris, K., Bullmann, T., Bakkum, D. J. & Frey, U. Revealing neuronal function through microelectrode array recordings. Frontiers in Neuroscience 8, 423 (2014). URL 10.3389/fnins.2014.00423.

[10] Liu, X., Gong, Y., Jiang, Z., Stevens, T. & Li, W. Flexible high-density microelectrode arrays for closed-loop brain–machine interfaces: a review. Frontiers in Neuroscience 18, 1348434 (2024). URL 10.3389/fnins.2024.1348434.

[11] Palanker, D., Mer, Y. L., Mohand-Said, S. & Sahel, J. Simultaneous perception of prosthetic and natural vision in amd patients. Nature Communications 13 (2022).

[12] Wang, B. et al. Electronic photoreceptors enable prosthetic vision with acuity matching the natural resolution in rats. Nature Communications 13 (2022).

[13] Lorach, H. et al. Photovoltaic restoration of sight with high visual acuity. Nature Medicine 21, 476–482 (2015).

[14] Ho, E. et al. Spatiotemporal characteristics of retinal response to network-mediated photovoltaic stimulation. Journal of Neurophysiology 119, 389–400 (2018). URL 10.1152/jn.00872.2016. PMID: 29046428.

[15] Ly, K. et al. Virtual human retina: Simulating neural signalling, degeneration, and responses to electrical stimulation. Brain Stimulation 18, 144–163 (2025). URL https://www.sciencedirect.com/science/article/pii/S1935861X25000154.

[16] Ly, K. et al. Residual photoreceptors affect the response of a degenerate retina to electrical stimulation. Proceedings of the National Academy of Sciences 123, e2537064123 (2026). URL https://www.pnas.org/doi/abs/10.1073/pnas.2537064123.

[17] Weiland, J. D. & Humayun, M. S. Retinal prosthesis. IEEE Transactions on Biomedical Engineering 61, 1412–1424 (2014).

[18] Beyeler, M. et al. A model of ganglion axon pathways accounts for percepts elicited by retinal implants. Scientific Reports 9, 9199 (2019).

[19] Ly, K. et al. Network-mediated subretinal stimulation is essential for prosthetic grating acuity. Investigative Ophthalmology & Visual Science 67, 4469 (2026).

[20] Chen, Z. et al. Photovoltaic implant simulator reveals resolution limits in subretinal prosthesis. Journal of Neural Engineering 19 (2022).

[21] Jensen, N., Ly, K., Kochnev Goldstein, A., Devaud, Q. & Palanker, D. Maximizing the fidelity of a photovoltaic subretinal prosthesis for human patients. Journal of Neural Engineering 23, 036034 (2026). URL 10.1088/1741-2552/ae6d77.

[22] Wilke, R. G. H., Khalili Moghadam, G., Lovell, N. H., Suaning, G. J. & Dokos, S. Electric crosstalk impairs spatial resolution of multi-electrode arrays in retinal implants. Journal of Neural Engineering 8, 046016 (2011). URL 10.1088/1741-2560/8/4/046016.

[23] Ozden, M. & Karagoz, I. Spatio-temporal electrode mapping and local interleaved stimulation method (STEMLIS) for artificial sight systems. Journal of Bioengineering & Biomedical Science 5, 1000169 (2015). URL 10.4172/2155-9538.1000169.

[24] Newman, J. Resistance for flow of current to a disk. Journal of the Electrochemical Society 113, 501 (1966).

[25] Palanker, D., Mer, Y. L., Mohand-Said, S., Muqit, M. & Sahel, J. Photovoltaic restoration of central vision in atrophic age-related macular degeneration. Ophthalmology 127, 1097–1104 (2020).

[26] Shin, A. et al. Amorphous silicon resistors enable smaller pixels in photovoltaic retinal prosthesis. Journal of Neural Engineering 22, 056017 (2025). URL 10.1088/1741-2552/ae0522.

[27] Boinagrov, D. et al. Photovoltaic pixels for neural stimulation: Circuit models and performance. IEEE Transactions on Biomedical Circuits and Systems 10, 85–97 (2016).

[28] Jensen, N., Chen, Z. C., Goldstein, A. K. & Palanker, D. Accelerated simulation of multi-electrode arrays using sparse and low-rank matrix techniques. IEEE Transactions on Biomedical Engineering 72, 2321–2330 (2025).

[29] Chen, Z., Ryzhik, L. & Palanker, D. Current distribution on capacitive electrode-electrolyte interfaces. Phys. Rev. Appl. 13, 014004 (2020). URL https://link.aps.org/doi/10.1103/PhysRevApplied.13.014004.

[30] Werginz, P., Wang, B.-Y., Chen, Z. C. & Palanker, D. On optimal coupling of the ‘electronic photoreceptors’ into the degenerate retina. Journal of Neural Engineering 17, 045008 (2020). URL 10.1088/1741-2552/aba0d2.

[31] Muqit, M. M. K., Mer, Y. L., Holz, F. G. & Sahel, J. A. Long-term observations of macular thickness after subretinal implantation of a photovoltaic prosthesis in patients with atrophic age-related macular degeneration. Journal of Neural Engineering 19, 055011 (2022). URL 10.1088/1741-2552/ac9645.

[32] Dommel, N. B. et al. A CMOS retinal neurostimulator capable of focussed, simultaneous stimulation. Journal of Neural Engineering 6, 035006 (2009). URL 10.1088/1741-2560/6/3/035006.

[33] Zwolan, T. et al. Comparison of continuous interleaved sampling and simultaneous analog stimulation speech processing strategies in newly implanted adults with a Clarion 1.2 cochlear implant. Otology & Neurotology 26, 455–465 (2005). URL 10.1097/01.mao.0000169794.76072.16.

[34] Loudin, J. D., Cogan, S. F., Mathieson, K., Sher, A. & Palanker, D. V. Photodiode circuits for retinal prostheses. IEEE Transactions on Biomedical Circuits and Systems 5, 468–480 (2011).

[35] Chen, Z. C., Wang, B.-Y. & Palanker, D. Harmonic-balance circuit analysis for electro-neural interfaces. Journal of Neural Engineering 17, 035001 (2020). URL 10.1088/1741-2552/ab89fd.

[36] Lorach, H. et al. Performance of photovoltaic arrays in-vivo and characteristics of prosthetic vision in animals with retinal degeneration. Vision Research 111, 142–148 (2015). URL 10.1016/j.visres.2014.09.007.

